# PREPRINT: Using digital epidemiology methods to monitor influenza-like illness in the Netherlands in real-time: the 2017-2018 season

**DOI:** 10.1101/440867

**Authors:** PP Schneider, CJAW van Gool, P Spreeuwenberg, M Hooiveld, GA Donker, DJ Barnett, J Paget

## Abstract

**Introduction:** Despite the early development of Google Flu Trends in 2009, digital epidemiology methods have not been adopted widely, with most research focusing on the USA. In this article we demonstrate the prediction of real-time trends in influenza-like illness (ILI) in the Netherlands using search engine query data.

**Methods:** We used flu-related search query data from Google Trends in combination with traditional surveillance data from 40 general sentinel practices to build our predictive models. We introduced an artificial 4-week delay in the use of GP data in the models, in order to test the predictive performance of the search engine data.

Simulating the weekly use of a prediction model across the 2017/2018 flu season we used lasso regression to fit 52 prediction models (one for each week) for weekly ILI incidence. We used rolling forecast cross-validation for lambda optimization in each model, minimizing the maximum absolute error.

**Results:** The models accurately predicted the number of ILI cases during the 2017/18 ILI epidemic in real time with a mean absolute error of 1.40 (per 10,000 population) and a maximum absolute error of 6.36. The model would also have identified the onset, peak, and end of the epidemic with reasonable accuracy

The number of predictors that were retained in the prediction models was small, ranging from 3 to 5, with a single keyword (‘Griep’ = ‘Flu’) having by far the most weight in all models.

**Discussion:** This study demonstrates the feasibility of accurate real-time ILI incidence predictions in the Netherlands using internet search query data. Digital ILI monitoring strategies may be useful in countries with poor surveillance systems, or for monitoring emergent diseases, including influenza pandemics. We hope that this transparent and accessible case study inspires and supports further developments in field of digital epidemiology in Europe and beyond.

## Introduction

Previous studies suggest that traditional disease surveillance systems could be complemented with information from online data sources.[1–3] The underlying premise is that people, nowadays, often turn to the internet when they face health problems.[4] With influenza-like illness (ILI), individuals might search for information about symptoms, look for remedies, or share messages on social media. All of these interactions leave digital footprints, which, when aggregated, could be harnessed to monitor disease activity.[1] In this way, online data streams could be used to support the timely detection of infectious disease outbreaks.

This hypothesis is not new, and in 2009 researchers at Google reported that their ‘Flu Trends’ model was able to predict ILI activity in the United States in real-time, by monitoring millions of queries on their search engine.[5] The aim of Google Flu Trends was to bridge a two-week lag in the reporting of ILI cases in the official surveillance statistics. Initially, the project indeed appeared to provide accurate predictions and was expanded to cover 29 countries around the world. In 2012, however, the model’s performance deteriorated, and in early 2013, it overestimated the peak of the epidemic by more than 140%. The failure, and subsequent termination of Google Flu Trends, received a lot of media attention and it also sparked an intense debate about the limitations of ‘big data’ in epidemiological research.[3,6]

Since then, the number of scholarly articles published in the field of digital epidemiology has grown significantly.[2,7] The discipline is, nevertheless, in an early stage and should still be considered as being experimental. Especially outside of the United States, there has been little effort to investigate the value of online data sources for epidemiological purposes.

Building on previous work,[8] our study assesses whether online search queries can be used to predict the ILI epidemic in the Netherlands during the 2017-2018 winter in real-time. Our investigation is meant to be an accessible case study in digital epidemiology. The full source code and data are provided under open license to encourage the application of this method to other countries and to other areas of epidemiological research.

## Methodology

### ILI-data

Weekly data on consultations for ILI were collected through sentinel practices participanting in the Nivel Primary Care Database,[9], and released by the European Centre for Disease Prevention and Control.[10] The practices constitute a nationally representative group of 40 general practices in the Netherlands. The methodology is further described by Donker.[11] It is important to note that the data are available in real-time and that the four-week reporting lag that we have assumed in this paper was only simulated for the purpose of our study.

### Web search queries

Data on web search queries were retrieved from Google Trends[12]. This online service provides statistics on how often a particular keyword was searched relative to the total search-volume across various regions of the world. It also provides information on related search queries (“people who searched for this, also searched for that”).

To identify potentially relevant keywords for our study, we retrieved the 25 terms that were most related to the search term ‘griep’, the Dutch word for ‘flu’. All keywords that contained a year were excluded, because they were - a priori - expected to be poor predictors of ILI rates in other years. Further a priori filtering of search terms wasnot performed. For the remaining search terms, we used the R-package gtrendsR[13] to download five years of weekly query statistics (from week 33/2013 to week 31/2018) for the Netherlands from Google Trends. Subsequently, we expanded the set of predictors by multiplying each predictor with all other predictors (to account for one-level interactions) and with themselves (to account for non-linearities).

### Modeling

We simulated the weekly use of a statistical model for predicting ILI incidence rates in real-time, based on Google search query statistics, during the 2017/18 ILI epidemic in the Netherlands. We assumed a four-week delay between the identification and the reporting of ILI cases, so that at week *w*, the ILI incidence of week *w* – 4 became available. For each week, the model was updated and re-fitted with the most recently available information (the search query data up until the current week), and the ILI incidence data up until four weeks ago. The updated model was then used to predict the ILI incidence for the current and the previous three weeks (i.e. to fill the assumed four-week reporting lag). To account for the temporal structure of the analysis, we set up an automated loop that iteratively splits the data into training and validation sets. The training data was used to fit a prediction model and the validation set was used to evaluate the model’s predictions on new data points.

### Automated analysis loop

The first four years of data (week 33/2013 to 30/2017) were used as training data only, while the analysis loop was run on the 52 weeks of the influenza season 2017/2018 (week 31/2017 to 31/2018). At each week, the data were split into a training set (for which both the ILI and Google data were made available to the model) and a four week validation set (for which only Google data were made available). This means, at the *ith* iteration of the loop, the analysis contained 207 + *i* weeks of training data (week 1 to 207 + *i*), and 4 weeks of validation data (week 207 + *i* + 1 to 207 + *i* + 4). Each week, a new model was built to predict the ILI incidence of the current week and the previous three weeks.

The model building process included the following steps. Firstly, dependent and independent variables were normalized and centered, with a mean of zero. This was performed separately for training and validation data, to prevent information leaking from the validation set. Variables with near zero variance were removed. We then used lasso (least absolute shrinkage and selection operator) regression,[14] in combination with cross validation (CV), to determine the optimal set of predictors and their regularized coefficients. See Technical Appendix (and the provided source code) for a detailed outline of the methods.

Lasso regression performs simultaneous variable selection and coefficient estimation. It imposes a penalty on the absolute values of the coefficients in the least squares estimation. In effect, less important parameters shrink towards zero, resulting in a selection of predictors, if coefficients become zero. The model’s complexity is controlled by the penalty parameter λ. Rolling forecast CV for time series, with fixed origin and expanding window, was used to find the optimal value for λ.

CV for time series is a variation of leave-k-out CV, which can be used to avoid the leakage of information from future to past observations. Similar to our automated analysis loop, CV for time series splits the data iteratively into a training set (the first *k* weeks) and a test or ‘hold-out’ set (the subsequent 4 weeks). In the first CV iteration, a lasso regression model is fit on data from week 1 to *k* = 52 and its predictions are tested on hold-out data from week 53 to 56. The process then ‘rolls’ forward, week-by-week, keeping the origin at week 1, and using an expanding number of weeks as training data with *k* = 52 + 1, +2,…, +*m*, whereby *m* increases with each iteration *i* of the outer analysis loop, with *m* = 207 − 52 + *i*. The prediction error over all *m* four-week hold-out sets is then aggregated to assess how well the statistical model can predict new data points. At each iteration of the outer analysis loop, the inner CV loop is run for 100 values of the penalty parameter λ, (ranging from 10^−8^ to 10^1/4^).

The λ_*i*_ of the model with the lowest maximum absolute error in the CV hold-out sets was used to fit the *ith* final lasso regression model on 207 + *i* weeks of training data, and to predict the ILI incidence for weeks 1-4 of the *ith* validation set. We used the *maximum* instead of the more commonly used *mean* absolute error as the selection criterion to increase model stability.

### Model evaluation

We analyzed the predictions of 52 lasso regression models (one for each week of year 5). Each model provided four predicted values, corresponding to the four weeks of the validation sets (except the first/last three models which had shorter horizons). We refer to the prediction of the current week as week 4 prediction (= 4 weeks since the last updated with official ILI data), and to the predicted values for the previous three weeks, as week 3 −1 predictions. Similarly, for all but the first three weeks of the validation period, there were four predicted values available.

We plotted the observed against the predicted ILI incidence values and assessed the performance of the statistical models over the validation period in terms of the mean absolute error (MAE). The accuracy of the week 1 - 4 predictions was evaluated separately and all values were back-transformed to their original scale. For comparison with previous studies, we also report the Pearson and Spearman correlation coefficients - however, correlations are not based on the absolute differences between the predicted and observed values and might, therefore, generate conflicting or misleading results. We also assessed how accurately the model predicted the onset and peak of the season. Furthermore, we investigated which search query terms were retained as predictors in the 52 models.

### Source code and data availability

The R-code for this study is provided under open CC BY license and the data that were used for this study can be accessed online.[15]

### RESULTS

#### Google search queries

We retrieved information on 26 search terms (‘Griep’ and the 25 most related keywords) from Google Trends (see table 1). Six terms were excluded from the analysis, as they contained a year. For all other terms (n=20), weekly search query statistics from 2013-08-12 to 2018-08-06 were downloaded. Overall, there was a high correlation between the query statistics, with an average correlation coefficient of 0.70 (minimum = 0.24; maximum = 0.97). Variables for first-order interactions between predictors (n = 190) and quadratic terms (n = 20) were included in the model. Together with the original keywords (n = 20), a total of 230 variables were considered in the analysis as potential predictors of the ILI incidence.

**Table 1:** Search terms retrieved from Google Trends

|  |  |  |
| --- | --- | --- |
| Griep symptomen griep symptomen de griep griep 2018* | tegen griep griep 2015* griep 2017* symptomen griep 2018* griep heerst | griep wat te doen griep zwanger ziek griep spierpijn hoofdpijn griep |
| griep koorts koorts griep 2016* griep hoe lang | verkoudheid hoe lang duurt griep heerst er griep griep 2014* | verschijnselen griep hoofdpijn griep nu |

\* Terms were removed from further analysis

#### Real-time ILI incidence prediction models

We simulated the weekly use of a real-time prediction model during the 52 weeks of the 2017/18 ILI season in the Netherlands. At each week, a new prediction model was built to estimate the ILI incidence of the current week (week 4) and the previous three weeks (weeks 3-1). Figure 1 shows the values of these week 1-4 predictions separately against the observed ILI incidence.

**Figure 1:**
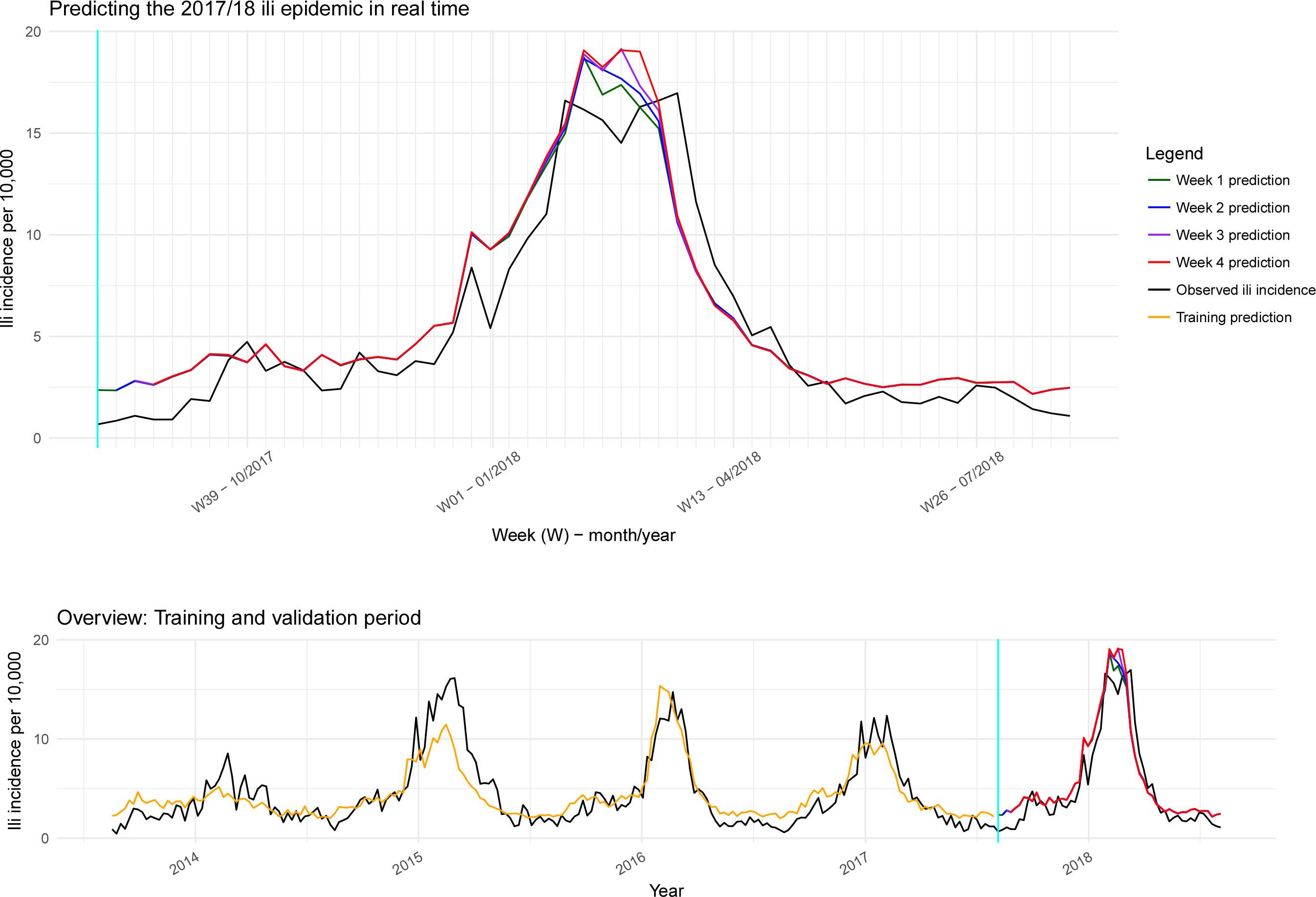
The time series plot shows the observed ILI incidence from week 16/2013 to week 15/2018 against the predictions of the 52 final lasso regression models. For the validation period (top), each model provides estimates for the week in which it, was ran (week 4) and the previous three weeks (week 3-1). In addition, predicted values within the training set, are provided for the first lasso regression model (bottom).

Overall, the 52 final lasso regression models predicted the ILI incidence with reasonable accuracy. The MAE for weeks 1,2,3, and 4 were 1.31, 1.35, 1.38, and 1.40. The corresponding Pearson correlation coefficients ranged between 0.95and 0.94, and Spearman coefficients were 0.90 for all four weeks.

Before the start of the epidemic in week 50/2017, the error was generally low, but it increased during the onset, and especially during the epidemic peak, when the highest prediction error (= 6.36) was registered (week 10/2018). After week 09-10/2018, the incidence was underpredicted by the models.

The model’s MAE in the validation period was slightly lower than the MAE observed in the CV hold-out sets (CV MAE for 1,2,3, and 4 predictions were 1.486, 1.547, 1.625, and 1.676). The maximum absolute error was markedly lower in the CV hold out sets (1.72).

The bottom plot in Figure 1 provides an overview of the entire five year observation period. The vertical blue line separates the training (left) from the validation period (right). For comparative purposes, the predicted values for the training period of the first prediction model (run in week 31/2017) are provided (first model training MAE = 1.32; maximum error 7.18). The figure also illustrates the seasonality of ILI epidemics (black line). The epidemic in 2017/18 had a slightly higher intensity and lasted longer than average, but was otherwise not exceptional.

#### Temporal aspects of ILI incidence predictions

From the visual presentation in Figure 1, it might be difficult to get assess what information was available at which week. To illustrate the temporal dynamics of the ILI prediction model, Figure 2 shows the results of 5 models at different points in time.

**Figure 2:**
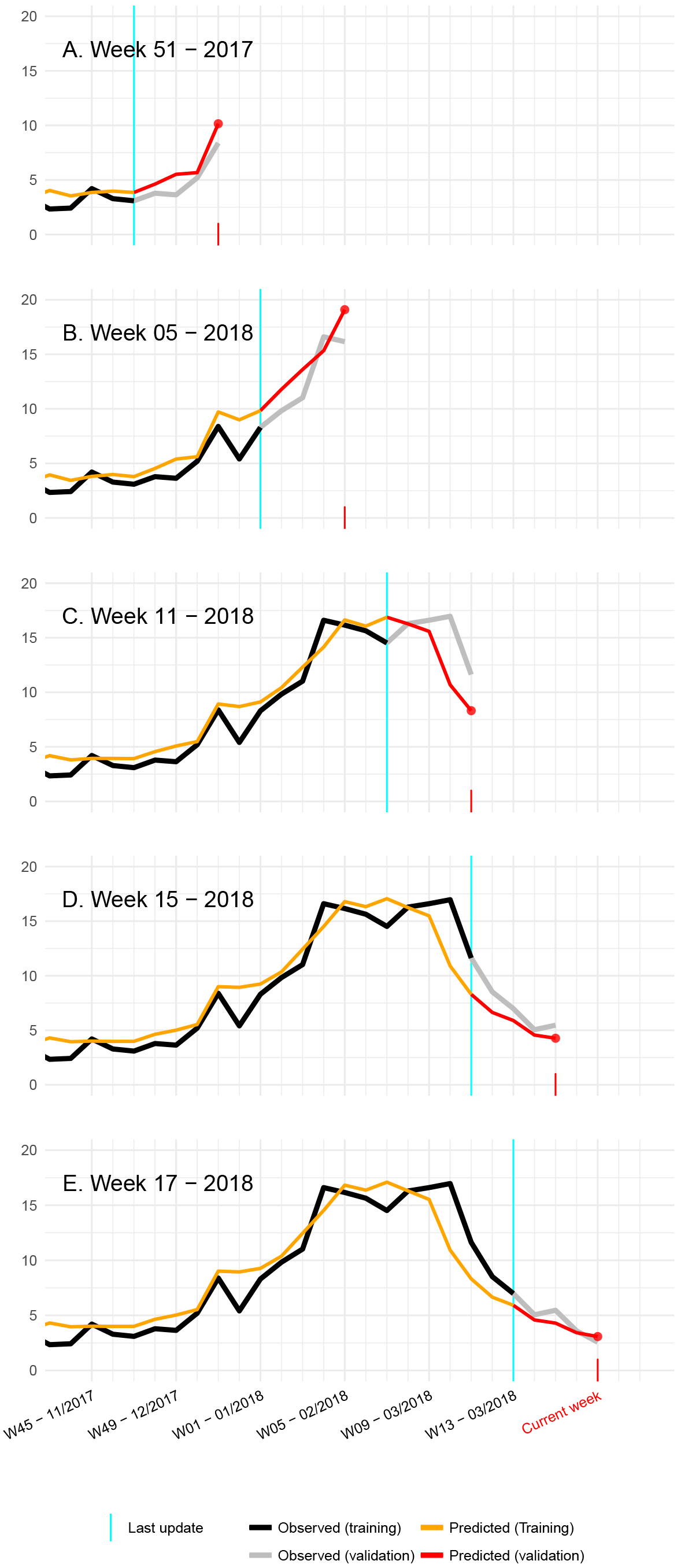
Observed versus predicted ILI incidence, showing data available at five points during the 2017/18 ILI season. The plots illustrate what information the prediction models would have provided at that particular point in time, if they were used. The y-axis shows the ILI incidence per 10,000 inhabitants; the week for which the last official ILI incidence data were made available is marked by the vertical cyan line; the current week, i.e. the week in which the model is used, is indicated by a red tick mark.

The model would have indicated the onset of the season one week ahead of the sentinel surveillance data (Panel A: observed onset = week 50/2017, predicted onset = week 49/2017). The peak of the season was predicted in week 07/2018 (Panel B), while the observed peak was biphasic with the highest incidence (=16.97) in week 10/2018 and the second highest (=16.6) in week 04/2018 (Panel C and D). The end of the season (i.e. ILI incidence falls below 5.1 per 10,000 for two consecutive weeks) was predicted in week 14/2018, and observed in week 16/2018 (Panel E).

Visual inspection indicates that the predictions generally appeared to be ahead of the actual ILI incidence. In fact, throughout the validation period, week 4 predictions were forecasting the ILI incidence of the coming week slightly more accurately (MAE = 1.11), than predicting the current week (MAE = 1.4).

#### Model specifications

Figure 3 provides an overview of the 52 weekly sets of predictors and their coefficients used in the final prediction models. During the validation period, the number of variables that were retained in the models as predictors ranged from 3 to 5. However, one predictor (‘Griep’ = ‘Flu’) had by far the most weight in all models, especially after week 32 of the validation period (= week 11 - 2018).

**Figure 3:**
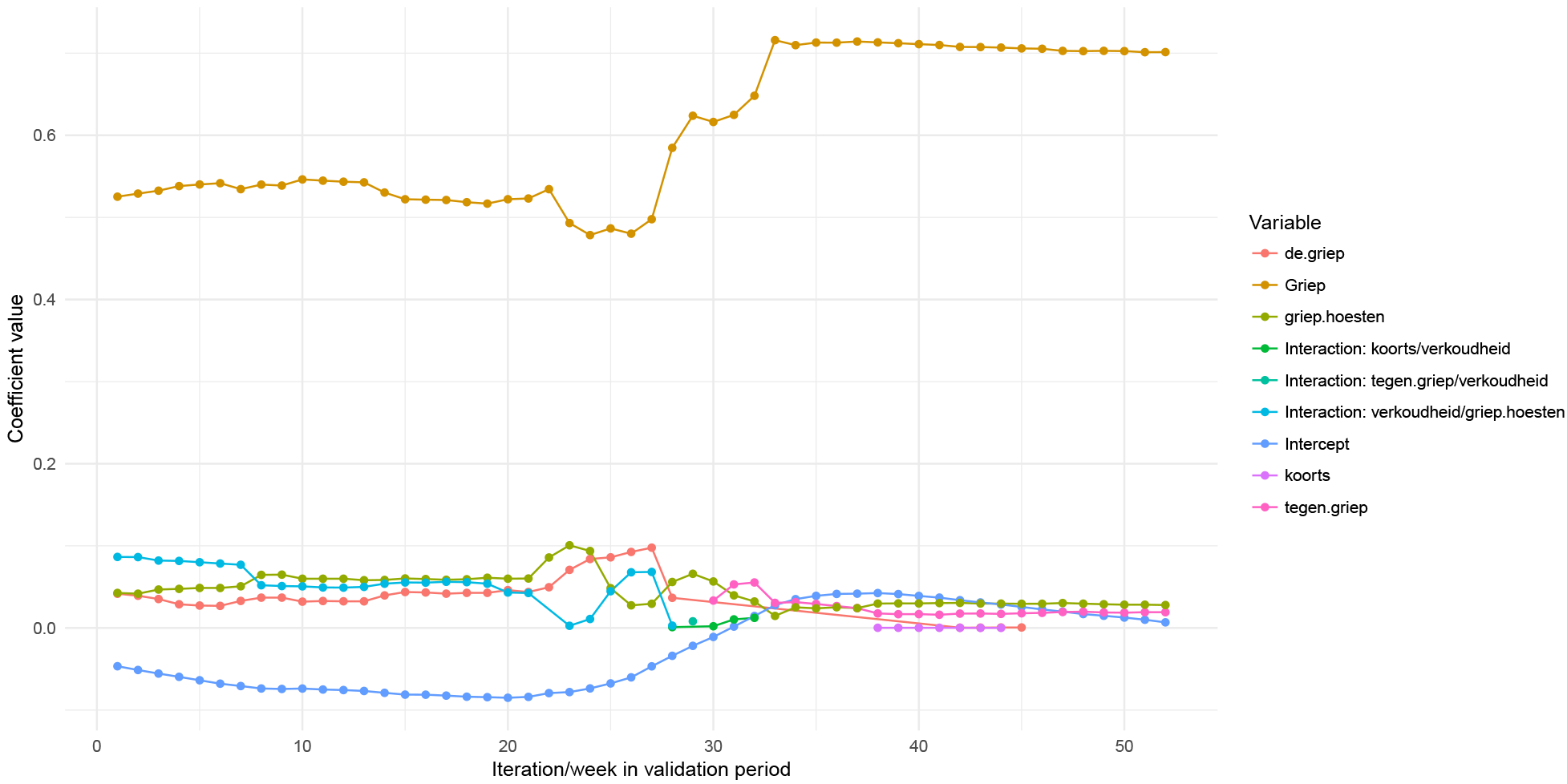
Predictors retained in the final lasso regression models throughout the 52 iterations. Coefficients with a value of zero are not shown.

### Discussion

Our study demonstrates that a statistical model, based on online search queries, can be used to predict the ILI incidence in the Netherlands during a normal influenza season. Assuming a four-week reporting lag, our model predicted the 2017/18 ILI epidemic in real-time with reasonable accuracy. The model would have identified the onset, peak, and end of the epidemic with one to three weeks difference.

This investigation provides an accessible but rigorous case study in digital epidemiology. The modelling steps are tractable and computationally economical, so that the source code can be modified and transferred to other settings. Indeed, we would encourage others to try this method in other countries and other infectious diseases or outcomes with a seasonal pattern (e.g. hayfever, asthma).

A feature of our study, that is worth highlighting, is the week-by-week simulation of the prediction model. We built 52 models, each of which was validated on four weeks of data (which were later used for fitting subsequent models). This iterative analysis loop allowed us to set a realistic framework for investigating our research question. Our results reflect how well a model would have performed, and what information it would have provided if it was used during the 2017/18 ILI season. The loop structure also enabled us to continuously update the model, as suggested by previous research[6,16], to prevent deterioration of performance. Each week, we re-fitted the prediction model using the most recently available ILI data from 4 weeks ago. We took this a step further and also repetitively applied CV to select the momentarily optimal set of predictors. Interestingly, most retained variables only changed marginally over time, and one predictor (‘Griep’) had by far the highest weight in all prediction models.

When preparing this project, we assessed a number of different data sources (e.g. Google searches and Wikipedia page visits),[8] but chose the data from Google Trends[12] to be the most advantageous for our project. The Google Trends service is publicly available, easy to use, and it covers the online search behavior of a majority of people in the Netherlands. It did come with certain limitations that should be considered when interpreting our findings or applying this methodology. Using the public API, weekly data can only be retrieved for periods of less than five years, which sets a boundary for the observation period. There is also a quota for the number of search requests, which limits the amount of predictor data that can be downloaded. Moreover, we cannot rule out that information was leaked from future to past observations, since we retrieved all data after the end of the season. If the data were retrieved each week during the season, results could have been different.[17] This is a point that should be assessed in the future.

In our modelling approach, we intentionally compromised on accuracy to provide a fast and comprehensible model. To further improve the predictive performance of our method, the model could be expanded by adopting strategies that have been successfully applied in other studies, including the use of more predictors (e.g. unrelated search queries and autoregressive terms)[18], additional data sources (e.g. Wikipedia page views, Twitter activity)[19,20], more extensive data pre-processing (e.g. principal component analysis), and alternative statistical models (e.g. ensemble methods)[7,20].

Our results are comparable to previous studies from European countries. Valdivia et al.[21] used historic data from the now terminated Google Flu Trends project and compared predicted ILI rates against sentinel surveillance estimates across the 13 European countries during the 2009/10 influenza pandemic. They found high correlations, with Spearman coefficients ranging from .72 in Poland to .94 in Germany. For the Netherlands, the authors report a correlation of .86, which is slightly lower than what we found in our study (= 0.90). More recently, Samaras et al.[22] studied the association between ILI incidence and influenza-related Google search queries in Greece and Italy in 2011 and 2012. They found Pearson correlation coefficients between .831 for Greece and .979 for Italy. It should be noted, however, that these figures were not validated on a test data set.

Numerous other studies, mostly from the US, have aimed to predict ILI incidence rates from online data. They have used various data sources and applied an array of different methods (Milinovich et al.[2] and O’Shea et al.[7] provide informative overviews). Unfortunately, many of the published studies suffer from methodological limitations, such as the use of inappropriate outcome measures (e.g. Spearman Rho), the absence of a rigorous validation method (e.g. using a single data set to fit a model and evaluate its predictions), or insufficient reporting (which does not allow for replication of results). Tabataba et al.[23] and Generous et al.[24] have published in-depth discussions of these points.

### Implications and outlook

In the Netherlands, there is no obvious case for monitoring ILI with these methods. ILI data, including virological information, are collected from sentinel practices on a weekly basis. However, during weeks 52/2017 and 01/2018, we made an interesting observation: the sentinel surveillance data indicated a temporary drop in the ILI incidence but we find the signal was likely not to have been caused by a decrease in the number of ILI cases, but rather by low health care utilization and/or changes in doctors’ working hours during the Christmas/New Year holiday period. For these two weeks, it could be argued that our prediction model could have usefully complemented the sentinel surveillance system.

Further potential applications of this digital epidemiology method include the provision of supportive low-cost online surveillance methods in resource-poor countries (e.g. countries which report data more slowly than in the Netherlands or do not cover all regions of a country) and an early warning system for pandemic outbreaks.[1] However, before these novel methods can be used in routine practice, more research is needed to better understand where, when, and how online surveillance can complement established systems in all world regions. The value of digital epidemiology still needs to be determined. To allow for incremental learning, future studies should report methods in sufficient detail, ideally providing the underlying source code and data.

## Conclusions

Our study demonstrates that a prediction model, based on online search queries, could have predicted the 2017/18 ILI epidemic in the Netherlands in real-time. The intensity of the epidemic, as well as its onset, peak, and end were estimated with reasonable accuracy. The value of using online surveillance methods to complement traditional disease surveillance systems should be further explored.

